# Infants’ neural oscillatory processing of theta-rate speech patterns exceeds adults’

**DOI:** 10.1101/108852

**Authors:** Victoria Leong, Elizabeth Byrne, Kaili Clackson, Naomi Harte, Sarah Lam, Kaya de Barbaro, Sam Wass

## Abstract

During their early years, infants use the temporal statistics of the speech signal to boot-strap language learning, but the neural mechanisms that facilitate this temporal analysis are poorly understood. In adults, neural oscillatory entrainment to the speech amplitude envelope has been proposed to be a mechanism for multi-time resolution analysis of adult-directed speech, with a focus on Theta (syllable) and low Gamma (phoneme) rates. However, it is not known whether developing infants perform multi-time oscillatory analysis of *infant*-directed speech with the same temporal focus. Here, we examined infants’ processing of the temporal structure of sung nursery rhymes, and compared their neural entrainment across multiple timescales with that of well-matched adults (their mothers). Typical infants and their mothers (N=58, median age 8.3 months) viewed videos of sung nursery rhymes while their neural activity at C3 and C4 was concurrently monitored using dual-electroencephalography (dual-EEG). The accuracy of infants’ and adults’ neural oscillatory entrainment to speech was compared by calculating their phase-locking values (PLVs) across the EEG-speech frequency spectrum. Infants showed better phase-locking than adults at Theta (~4.5 Hz)and Alpha (~9.3 Hz) rates, corresponding to rhyme and phoneme patterns in our stimuli. Infant entrainment levels matched adults’ for syllables and prosodic stress patterns (Delta,~1-2 Hz). By contrast, infants were less accurate than adults at tracking slow (~0.5 Hz) phrasal patterns. Therefore, compared to adults, language-learning infants’ temporal parsing of the speech signal shows highest relative acuity at Theta-Alpha rates. This temporal focus could support the accurate encoding of syllable and rhyme patterns during infants’ sensitive period for phonetic and phonotactic learning. Therefore, oscillatory entrainment could be one neural mechanism that supports early bootstrapping of language learning from infant-directed speech (such as nursery rhymes).

## 1 INTRODUCTION

As a first step in speech perception, the brain must parse the continuously varying energetic patterns of the speech signal into discrete phonological units (e.g. phonemes, syllables and stress patterns) that transmit meaning in one’s native language. Current neural models of adult speech processing suggest that the brain performs this segmentation or ‘chunking’ process through speech-brain *phase-locking* (Giraud & Poeppel, 2012; Ghitza, 2011; Gross et al, 2013) in which neuronal oscillations in the auditory cortex at different timescales *entrain* (phase-lock) to modulation patterns in the speech amplitude envelope on equivalent timescales. For adults, speech-brain entrainment is proposed to occur most strongly at the Theta rate (3-7 Hz), which coincides with the most dominant rhythmic rate in the adult speech modulation spectrum, and corresponds to syllable-rate patterns of utterance (~5 Hz, Greenberg et al, 2003; Luo & Poeppel, 2007). In parallel, faster oscillations in the low Gamma range (25-35 Hz) are thought to track quickly-varying phonetic information, such as formant transitions and voice-onset times, whilst slower Delta oscillations (1-3 Hz) could track stress and intonational patterns (Ghitza & Greenberg, 2009). Therefore, adults can concurrently track temporal patterns in speech across a range of different rates, but Theta-rate oscillators have been proposed to occupy a privileged role as “master oscillators” (Giraud & Poeppel, 2012; Ghitza, 2011), referring to their hierarchical modulation of faster-rate oscillations (such as Gamma) through phase-amplitude coupling. This Theta-driven process is biologically expedient given the rhythmic dominance of the syllable rate in the adult speech signal. However, it is not known if multi-timescale oscillatory speech parsing also occurs early in development during infancy, and if so, whether infants’ and adults’ profile of neural entrainment differs across speech timescales

### 1.1 A potential role for oscillatory entrainment in early language development

Neural entrainment, or phase-locking, is an efficient biological mechanism for temporal parsing of speech patterns because the phase of cortical oscillations reflects the excitability (receptiveness) of the underlying neural coding populations (Schroeder et al, 2008). Thus, speech information that has accumulated within the same oscillatory cycle is bound together (i.e. falls within the same receptive period), whereas speech information that straddles different oscillatory cycles is segregated into separate units - i.e. segmentation. However, if there are no landmarks in the speech signal to guide or constrain this oscillatory parsing mechanism, speech patterns can easily be mis-segmented. This is why it is crucial that both bottom-up stimulus-dependent mechanisms (e.g. perceptual saliency) and top-down control mechanisms (e.g. attention) act in concert to continuously re-align the phase of neural oscillations with their respective acoustic templates in the speech signal (Schroeder et al, 2008), thereby achieving accurate parsing of speech into its phonological constituents. This suggests that the fidelity of speech-brain phase-locking should be directly related to the accuracy of speech comprehension, and this has indeed shown to be the case in adults. For example, Ahissar and colleagues found that, when presenting compressed speech to adults, the comprehensibility of the speech signal correlated with the phase-locking observed between the temporal envelope of the stimulus and the subject’s cortical activity (Ahissar et al., 2001). Similarly, Luo & Poeppel, (2007) reported that the phase pattern of adults’ Theta (4–8 Hz) responses reliably discriminated between different sentences, and was also correlated with speech intelligibility (see also Peelle et al, 2013).

As neural entrainment studies have primarily been conducted with adults, it is not known whether infants also employ multi-time resolution oscillatory analysis of the speech signal with a Theta-rate focus, and if (and how) these mechanisms could potentially support language acquisition by infants. One key developmental milestone for language learning occurs by 10-12 months of age. By this age, ‘perceptual tuning’ to native phonetic categories occurs, so that infants’ ability to discriminate between non-native phonetic contrasts declines (Werker & Tees, 1984) whilst their sensitivity to native consonant sounds increases (Kuhl et al, 2008). Further, at 9 months, infants begin to show the ability to distinguish between unfamiliar words that either follow or violate the phonetic and phonotactic patterns of their native language (Jusczyk et al, 1993). These dramatic changes in infants’ speech processing abilities are thought to be driven by an intense phase of phonetic learning (Kuhl et al, 2008). During this sensitive or critical period for language learning, exposure to prosodically- and phonetically-enhanced speech (such as infant-directed speech, IDS) causes neural changes at the molecular level that attune the brain to the statistical and perceptual properties of one’s native language (Kuhl et al, 2008; Werker & Hensch, 2015).

Previous studies have already suggested that oscillatory mechanisms may be important for supporting several aspects of language development, such as neural commitment to native phoneme sounds (Ortiz-Mantilla et al, 2013; Bosseler et al, 2013) and also to native prosodic rhythm patterns (Pena et al, 2010). For example, Ortiz-Mantilla et al (2013) observed that the power of gamma oscillations in 6-month-old infants was selectively enhanced during the processing of native phoneme contrasts, but not non-native contrasts, suggesting that gamma oscillatory mechanisms could be implicated in infants’ learning of native phoneme sounds. Further, oscillatory entrainment at slower timescales (e.g. Theta, Delta) could support infants’ known ability to use more slowly-varying syllable and prosodic stress patterns in speech to boot-strap early language acquisition (Gleitman & Wanner, 1982; Morgan & Demuth, 1996; Leong & Goswami, 2015). For example, in a statistical learning task, Kabdebon et al (2015) reported that 8 month-old infant’s entrainment to syllable (Theta) rate patterns during the learning period predicted their later neural discrimination between rule-words and part-words at test. Therefore, it is of broad scientific relevance to investigate how the various timescales for oscillatory processing might support different aspects of language learning during early life.

### 1.2 Maturation of auditory processing mechanisms during early life

Convergent research suggests that even young infants can entrain to sounds with high temporal precision. One study using the auditory brainstem response found no significant difference in the temporal resolution of the auditory systems of 3-month-old infants versus adults, when measuring responses to brief gaps in noise (Werner et al, 2001). Using a combined EEG/NIRS approach, Telkemeyer et al (2009, 2011) studied sensitivity to temporally-modulated noise in neonates as well as in 3-month versus 6 month-old infants. They found that neonates could discriminate between different rates of temporal modulation (Delta to Gamma timescales) in non-speech sounds suggesting that the neural mechanisms for speech-brain entrainment may already be functional at birth. Further, research using the auditory steady state response (ASSR) suggests that even neonates show accurate neural entrainment to temporal rhythmic patterns in non-speech auditory stimuli (Picton et al, 1987; Rickards et al, 1994). Of note, infant ASSRs are significantly smaller than adults’ (between 1/3-1/2 adult amplitude; Lins et al, 1996), and thus are measured with a poorer signal-to-noise ratio, making it difficult to directly compare the fidelity of infant versus adult neural responses.

However, not all aspects of auditory processing mature early in life, in particular, the processing of complex speech sounds. Johnson and colleagues measured brainstem responses evoked by both click and speech syllables in 3-12-year-old children. They found that, whereas all children exhibited identical neural activity to a click, 3- to 4-year-old children displayed delayed and less synchronous onset and sustained neural response activity when elicited by speech compared with 5- to 12-year-olds (Johnson et al., 2008). Further, although cortical auditory evoked potential components (P1, N1, P2, and N2) can be recorded in prematurely-born infants as early as at 24 weeks gestational age, these waveforms do not show adult-like morphology and latency until late childhood (see Wunderlich & Cone-Wesson, 2006 for review). Other aspects of complex auditory processing, such as perception of speech in noise, continue to develop even into adolescence (see Moore, 2002 for review). Therefore, whilst basic neural entrainment mechanisms may be intact even from birth to support a preliminary temporal parsing of the speech signal into its fundamental phonological constituents, the neural mechanisms that support more complex processing of speech (e.g. to support comprehension, auditory scene analysis and robustness to noise) may be much slower to emerge over development.

### 1.3 Differences in temporal structure between infant-directed speech (IDS) and adult-directed speech (ADS) and possible effects on oscillatory entrainment

When studying speech-brain entrainment in infants, it is important to recognise that the speech that is commonly heard by infants differs markedly in temporal structure from the speech that is commonly heard by adults. Logically therefore, the temporal *profile* of neural entrainment to IDS should differ from ADS. When speaking to infants, adults spontaneously enrich the melodic and rhythmic patterning of their utterances so that infant-directed speech (IDS) is higher-pitched, slower and louder than adult-directed speech (ADS) (Fernald & Simon, 1989). These adaptations serve to capture infants’ attention and communicate affective warmth, and also support language learning directly (Fernald, 1989; Cooper & Aslin, 1989; Liu et al, 2003), as evidenced by the fact that infants favour listening to IDS over ADS (Fernald, 1985), and the infant brain shows enhanced processing of IDS-like formant-exaggerated speech sounds (Zhang et al, 2011). However, aside from these affective differences, the different *rhythmic* signature of IDS may also moderate neural temporal processing of speech, by heightening speech-brain entrainment at a different rhythmic rate for infants.

IDS is slower and contains more prosodic stress marking than ADS (Leong et al, in press), therefore, its modulation spectrum is shifted towards slower rates, with the strongest amplitude modulation and rhythmic regularity at the Delta rate, rather than at the Theta rate, which is the peak modulation rate for ADS (Leong et al, 2014; Greenberg et al, 2003). Since even neonates can utilise slow prosodic rhythm patterns to distinguish between languages of different rhythm typologies (Nazzi et al, 1998), this suggests that human infants are born with an already well-developed ability to detect prosodic rhythm patterns (that typically vary at the Delta rate). Thus, the rate difference between IDS and ADS presents an opportunity to test two interesting and contrasting predictions about the relationship between entrainment in the brain and speech temporal structure. If neural entrainment is entirely dependent on the temporal structure of the input auditory signal, then during infancy, Delta may play the role of "master oscillator" instead of Theta, since Delta is more dominant in the temporal structure of the IDS signal that must be parsed. Conversely, if Theta-rate oscillators are neurobiologically given to the oscillatory management of speech structure parsing, then infants should show enhanced processing at the Theta-rate even when parsing IDS (whilst still proficiently parsing structure at a Delta rate). Accordingly, here we assess and directly compare infants’ fidelity of neural entrainment relative to adults’ across the entire modulation spectrum of infant-directed speech (0.5 - 40 Hz), with a particular interest in their patterns of entrainment at Delta and Theta rates.

### 1.4 Experimental rationale, predictions and considerations

The current experiment was conducted to examine infants’ processing of the temporal structure of sung nursery rhymes, and to compare their neural entrainment (across multiple timescales) with that of well-matched adults (their mothers). We predicted that infants would show enhanced phase-locking (relative to adults) at the Theta rate when listening to IDS. To test this, we systematically assessed infants’ neural entrainment to the temporal modulation patterns of sung IDS across a wide range of temporal rates (0.5 Hz - 40 Hz) at two recording sites – C3, and C4. In particular, we chose to include very slow (e.g. 0.5 Hz) temporal rates in our analysis. Rates under 1 Hz are not typically examined in *adult* models of speech processing which focus on Delta- to Gamma–rate EEG frequency bands (Giraud & Poeppel, 2012; Ghitza, 2011). However, speech amplitude modulations as slow as 0.4 Hz also contribute toward speech intelligibility (Fullgrabe et al, 2009), and the IDS signal is rich in these slower modulation rates that carry phrasal and syntactic structure, as evidenced in our current stimulus material.

To reduce the amount of random variation between adult and infant groups, this study uniquely used *mothers* as the adult group. Mothers are genetically related to their infants and also share the same home language environment. We tested mother and infant concurrently in the same experimental session to reduce unwanted variations in stimulus presentation and to ensure consistency between their electrophysiological recordings. We then computed the degree of speech-brain entrainment observed in infants and in their mothers using the *Phase-Locking Value* (PLV; Lachaux et al, 1999), which measures phase synchronisation between speech modulation patterns and the EEG response. In addition, in the Supplementary Materials, we present equivalent analyses based on another commonly used measure, Coherence (Torrence & Compo, 1998; Grinsted et al, 2004), which is sensitive to signal amplitude in addition to phase. Given that infants’ EEG responses are often larger in amplitude than adults’ responses (Kurtzberg et al, 1984; Wunderlich et al, 2006), and infants’ EEG power spectra show greater power at low frequencies and less power at high frequencies than adults’ (see Figure S3 in Supplementary materials), speech-brain entrainment indices that are sensitive to amplitude (such as Coherence) will be biased by underlying differences in the adult and infant power spectra. For example, if infants show higher Coherence with speech at low frequencies as compared to adults, it will be unclear whether this result is due to their higher endogenous Delta power, or whether they are truly more proficient at tracking Delta-rate temporal patterns in the speech envelope. Consequently, a pure phase-locking measure such as the PLV that is insensitive to amplitude differences allows for a more straightforward interpretation of any observed differences between adults and infants.

## 2 METHODS

### 2.1 Participants

58 participants (29 mothers, 29 infants) participated in the study. The infants showed a 15M/14F gender split. All mothers were native English speakers and infants were aged between 6.3-14.8 months (median = 8.3 months, SD = 2.3 months). All infants had no neurological problems and had normal hearing and vision, as assessed by their mother’s report. Due to the large age range in our sample, we also assessed whether infants’ entrainment to the speech signal varied as a function of their age. There was no significant correlation between infants’ age at test and their phase-locking values (p>.05 for all comparisons, Bonferroni-corrected p-value threshold = .00179). Accordingly, infant age was not considered to be a confounding factor in the subsequent analysis.

### 2.2 Materials

The stimuli used were video clips of a female actress who sang familiar nursery rhymes, including ‘Twinkle Twinkle Little Star’ and ‘The Wheels on the Bus’. Prior to starting the experimental session, mothers confirmed that the nursery rhymes were familiar to both themselves and their infant. Seven different nursery rhyme clips were used in total, and each clip was repeated twice in a counterbalanced order across participants. The actress was filmed against a black background, and the camera angle showed the actress’ face but not her body, as shown in Figure 1.

**Fig 1.**
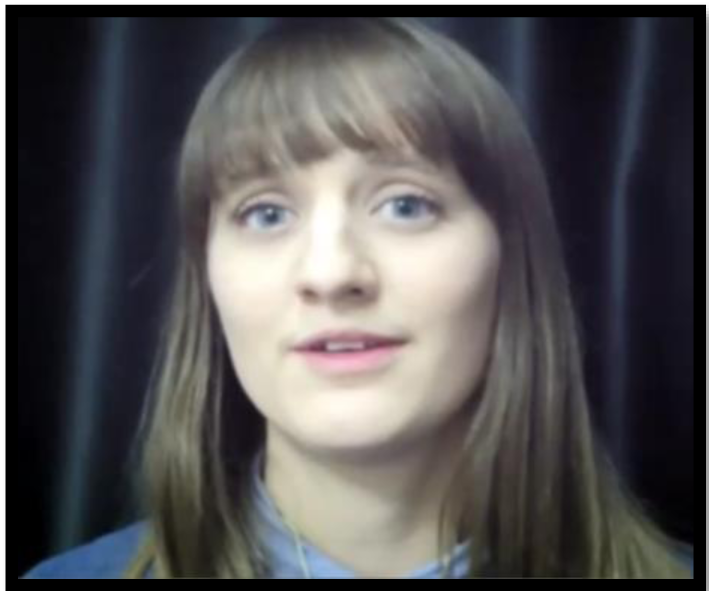
Screenshot of nursery rhyme video stimulus

The average duration of each nursery rhyme clip was 15.52 seconds (range 10.28s-23.69s) and the total duration of the nursery rhyme clips (with repetition) was 183s. The temporal rhythmic characteristics of the nursery rhyme stimuli are summarised in Table 1. Note that as the nursery rhymes were sung and not spoken, the stimuli contained longer pauses and greater vowel lengthening than non-sung speech, hence the temporal rates reported here are slower than for spoken infant-directed speech. For example, Fernald & Simon (1984), reported a spoken IDS syllable rate of 4.2 Hz, here the sung syllable rate was 2.61 Hz.

**Table 1.** Temporal rhythmic characteristics of the nursery rhyme stimuli

|  | Phrase rate | Stress foot | Syllable | Onset-Rime | Phoneme |
| --- | --- | --- | --- | --- | --- |

|  |  | (pulse) rate | rate | rate | rate |
| --- | --- | --- | --- | --- | --- |
| **Example of linguistic unit** | *"The clock struck one"* | *"The clock"* | *"clock"* | *"cl" (onset)*<br>*"ock" (rime)* | /k/ |
| **Mean (SD)** | 0.64 Hz<br>(0.13 Hz) | 1.28 Hz<br>(0.25 Hz) | 2.61 Hz<br>(0.54 Hz) | 4.99 Hz<br>(0.90 Hz) | 7.05 Hz<br>(1.31 Hz) |
| **Range** | 0.51 - 0.80<br>Hz | 1.01 - 1.60<br>Hz | 1.77 - 3.26<br>Hz | 3.50 - 6.16<br>Hz | 4.85 - 8.15<br>Hz |

The mean modulation spectrum (power spectrum of the amplitude envelope) of the 7 nursery rhyme stimuli is shown in Figure 2. The power profile of the rhymes followed a decreasing pattern, with the highest modulation power at low (Delta) frequencies, and the lowest modulation power at high frequencies. Note that this modulation spectrum for sung IDS differs from the typical modulation spectrum of ADS, which commonly shows a peak in modulation power in the Theta range (Greenberg et al, 2003).

**Figure 2.**
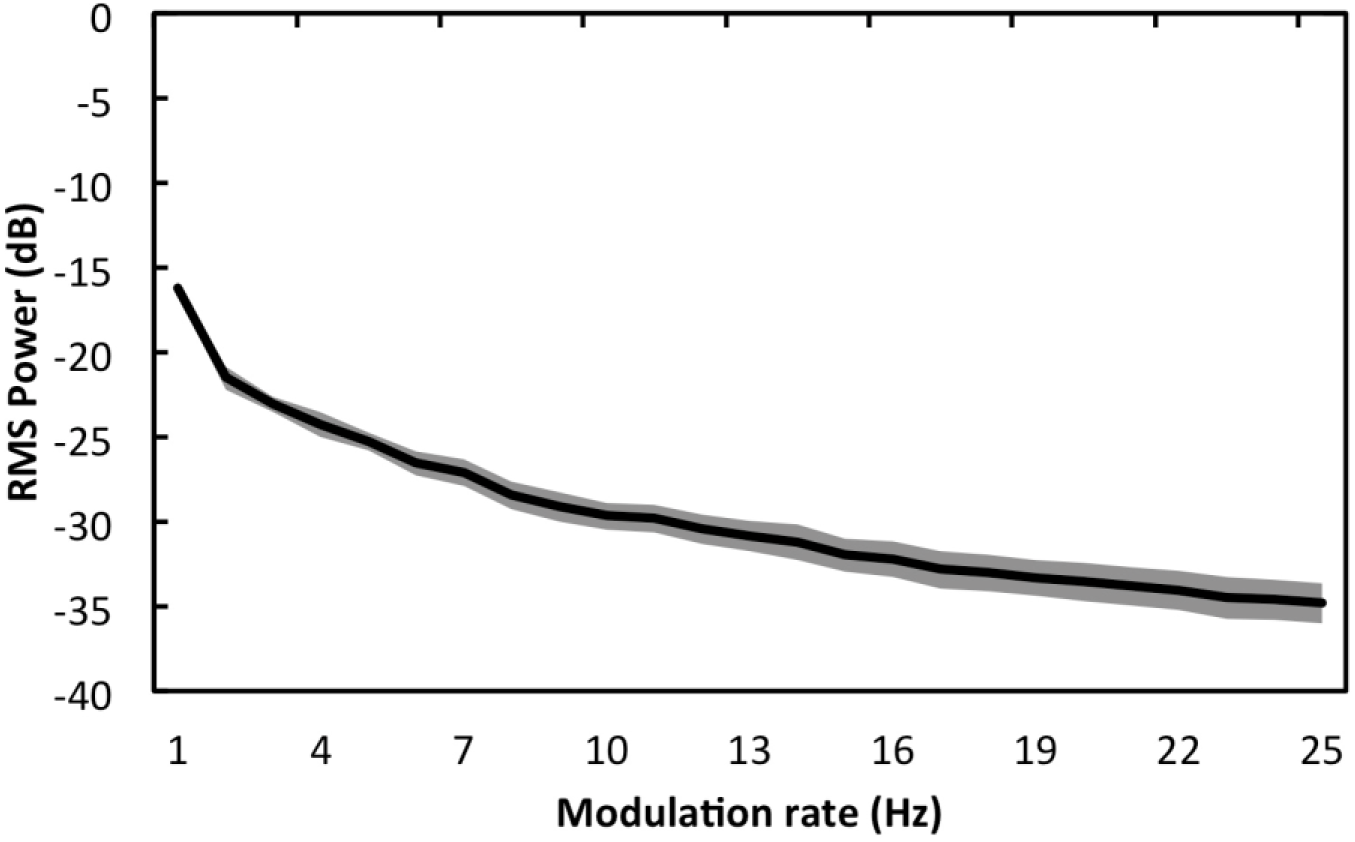
Mean modulation spectrum of 7 nursery rhyme stimuli. Shaded areas show the standard deviation observed between the different rhymes.

### 2.3 Protocol

Mothers and their infants were seated side by side, with infants sitting upright in a high chair, so that both faced the display screen throughout the video. The video clips were presented on a monitor subtending approximately 40° of visual angle, against black surrounds, and the sound was played through the speakers on the monitor at a level that was comfortable for infants (<75 dB SPL). To maximise infants’ attention on the video stimuli, an attention-getting sound was played prior to each video clip. To synchronise the EEG recording with the video stimuli, triggers were sent from the stimulus computer to the EEG acquisition software at the start of each video clip. Triggers were sent using a National Instruments USB 6501 device with latencies of ~1ms^1^. Only neural activity during the presentation of the videos was analysed. Mothers were instructed to watch and listen carefully to the video, and did so throughout. Infants were occasionally inattentive during stimulus presentation. Inattentive episodes were detected via video monitoring and coding (see Section 2.5 for further details).

### 2.4 EEG acquisition

EEG was recorded simultaneously from infants and their mothers from the central region (C3 and C4) and referenced to the vertex (Cz) according to the International 10–20 placement system. EEG was recorded from central sites to reduce potential confounding influences of muscle artefacts and blinking while still capturing a robust auditory response (the ASSR typically has a frontocentral topography; e.g. Saupe et al, 2009; Shuai & Elhilali, 2014). The vertex reference location was used because it produces comparable results to other reference sites (Tomarken, Davidson, Wheeler, & Kinney, 1992), and is the least invasive for young infants. Prior to electrode attachment, electrode sites were marked and wiped with alcohol. Electrodes were then affixed to the scalp using Signa conductive electrode gel (Parker Laboratories Inc, NJ). After the electrodes were in place, a soft elastic band was wrapped around the infant’s head to secure the electrodes. EEG signals were obtained using a Biopac MP150 Acquisition System with filters set at 0.1 Hz high pass and 100 Hz low pass. Wireless dual-channel BioNomadix amplifiers were used, which, due to the lack of tethering wires, reduced distraction for the infant during the experiment. EEG signals were recorded at 1000 Hz using AcqKnowledge software (Biopac Systems Inc). All further analysis was performed using Matlab software (The Mathworks Inc).

### 2.5 EEG selection and artifact rejection

To ensure that the EEG data used for analysis reflected only awake, attentive and movement artifact-free behaviour we performed a two-step selection and artifact rejection procedure. First, each mother-infant dyad was video-taped during the experimental session and these videos were reviewed frame-by-frame (frame rate of 30 fps) to identify the exact onset and offset times of all the infants’ movement artifacts, including blinks, head and limb motion, and chewing. Only time periods when infants were still and also looking directly at the nursery rhyme video were accepted. Next, manual artifact rejection was performed on this awake, attentive data to further exclude segments where the amplitude of infants’ or mothers’ EEG exceeded +100 μV (e.g. mechanical artifacts from wire motion). This two-stage artifact rejection procedure ensured that data were free of artefact due both to inattentiveness and other behavioural factors, and due to mechanical factors.

### 2.6 Speech-brain entrainment analysis : Phase locking value (PLV)

The EEG data was first low-pass filtered under 45 Hz using an inverse fft filter to remove line noise (EEGLAB eegfiltfft.m function; Delorme and Makeig, 2004). Next, the wholeband amplitude envelope of the speech signal of the video stimulus was extracted using the Hilbert transform. Frequency decomposition was performed by applying a continuous wavelet transform to the neural data and the speech envelope, which convolves each time series with scaled and translated versions of a wavelet function (Mallat, 1999). Here, the wavelet function chosen was the complex Morlet wavelet (bandwidth of mother wavelet = 1 Hz, time resolution = 0.1 Hz). The wavelet time-frequency decomposition was performed at 7 log-spaced frequencies as follows: 0.50 Hz (Slow), 1.03 Hz (Delta), 2.15 Hz (Delta), 4.47 Hz (Theta), 9.28 Hz (Alpha), 19.27 Hz (Beta) and 40.00 Hz (Gamma). The phase series at each frequency was extracted from the complex wavelet coefficients, and divided into matching EEG and speech epochs of length 2.0s (with an overlap of 1.0s). To assess the degree of entrainment between the neural EEG signal and the speech amplitude envelope the phase-locking value (PLV) was computed.

Following Lachaux et al (1999), one PLV estimate for each epoch at time window, *t*, was computed as follows:

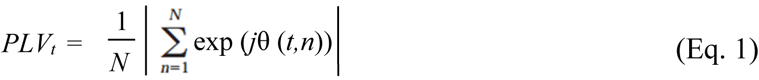

Where *N* is the number of data samples within the epoch, and θ *(t, n)* is the instantaneous phase difference between the EEG signal, *x*, and the speech envelope, *y* for each sample, *n,* within the epoch at time window, *t*:

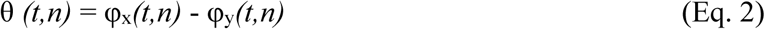

The PLV takes values between [0, 1], where a value of 0 reflects the absence of phase synchrony and a value of 1 reflects perfect synchronisation. The average PLV over all epochs for each participant was used for subsequent statistical analyses.

### 2.7 Methodological control analysis with white noise surrogates

To assess whether the data yielded levels of phase-locking that were higher than that which would be expected to occur by chance, a methodological control analysis was conducted with white noise surrogates. For every data sample from each participant, a white noise surrogate was generated by creating a random data series of the same length as the original sample. These surrogate data were then passed through the same analysis pipeline as the real data samples, and the mean levels of phase-locking observed were computed at each frequency. These control data means were then used as threshold values against which the actual data samples were tested using a series of one-sample t-tests. All the experimental data differed significantly from the white noise threshold values at all frequencies (p<.05 for all comparisons, corrected using the Benjamini-Hochberg False Discovery Rate procedure [Benjamini & Hochberg, 1995]), indicating that the levels of phase-locking observed in our data were above significantly above chance. For reference, these white noise thresholds are annotated in Figure 3 of the Results Section.

**Figure 3.**
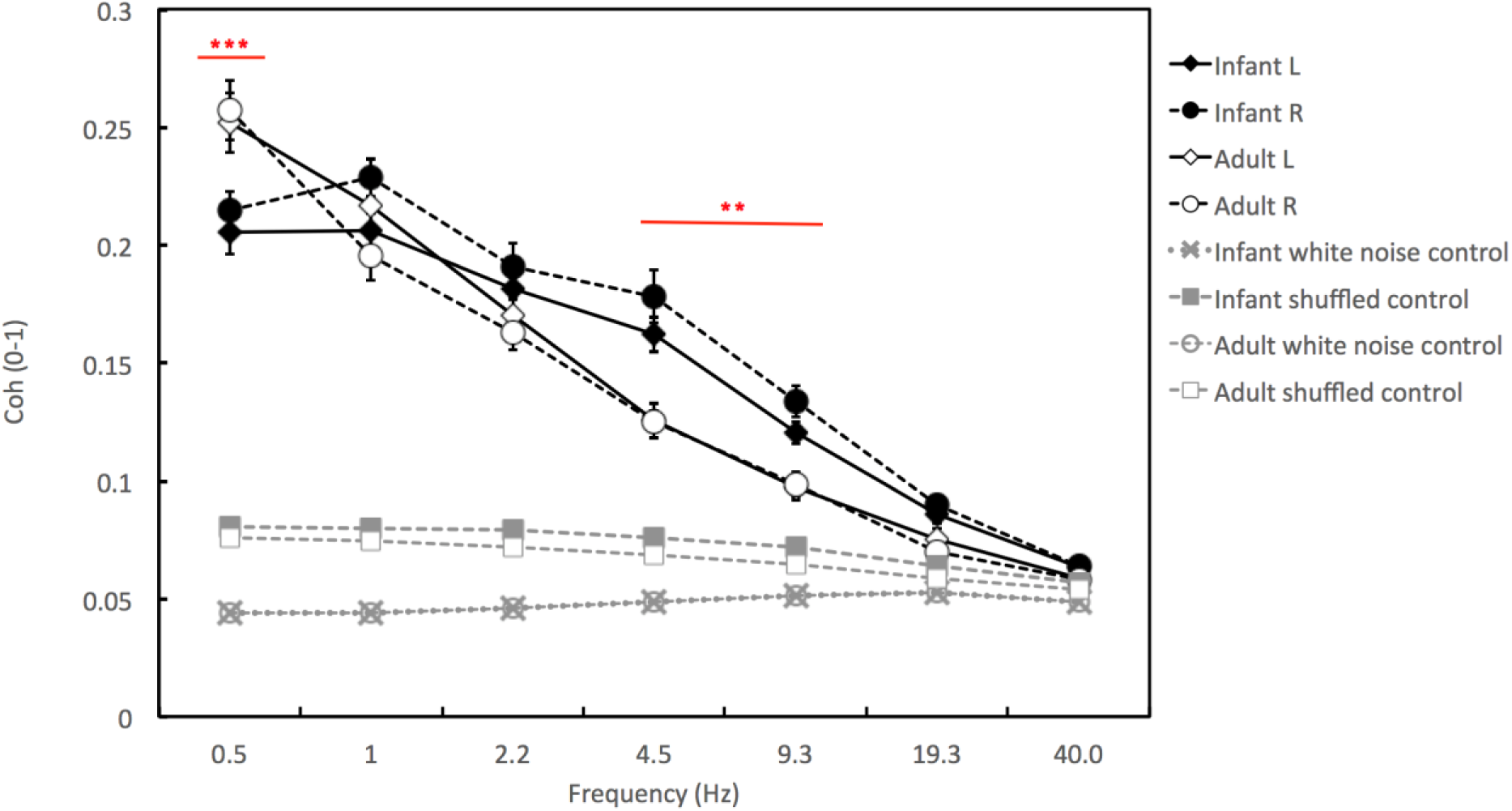
Phase-locking values (PLV) for infants and adults over left and right hemispheres, for 0.5-40 Hz. ‘Infant L’ and ‘Infant R’ show the results obtained over the left and right hemisphere electrodes, accordingly. ‘White noise control’ shows the results of the first control analysis conducted with white noise surrogates, as described in the Methods section. ‘Shuffled control’ shows the results of the second control analysis conducted with shuffled data, as described in the Methods section. Error bars show Standard Error of the Means. Stars indicate the significance of the adult vs infant Tukey post hoc HSD tests, conducted as described in the main text. ** *p<.01*, *** *p<.001*. Comparisons with the control data were significant for all comparisons with the exception of the Adult L vs Adult shuffled control comparison at 40 Hz only.

### 2.8 Methodological control analysis with temporally-shuffled data

To assess the potential effects of differences between infants’ and adults’ EEG dynamics on their measured PLV values, we conducted a second control analysis in which the EEG data were randomly shuffled (translocated) in time to destroy temporal correspondences between the EEG signal and the speech amplitude envelope. To ensure that even fine-grained temporal correspondences at the highest frequency measured (here 40 Hz) were removed, 20 ms-long segments (the period of a 50 Hz) cycle were randomly shuffled, therefore ensuring that temporal correspondences at all frequencies below 50 Hz were decimated. The same PLV metrics were then computed on this shuffled data. The resulting shuffled data PLV means were then used as threshold values against which the actual data samples were tested using a series of one-sample t-tests, for each frequency and hemisphere. For the infant data, all the experimental data differed significantly from the shuffled data values at all frequencies (p<.05 for all comparisons, corrected using the Benjamini-Hochberg False Discovery Rate procedure [Benjamini & Hochberg, 1995]). However, for the adult data, phase-locking at all frequencies except for the Gamma (40 Hz) rate over the left hemisphere was significantly higher than the shuffled data values (p<.05, corrected using the Benjamini-Hochberg False Discovery Rate procedure [Benjamini & Hochberg, 1995]). For reference, these shuffled data thresholds are also annotated in Figure 3 of the Results Section. It is interesting to note that the shuffled data values for infants were uniformly higher than adults’ across all frequencies, which could suggest that infants’ oscillations are more strongly reset by rhythmic patterns in speech. However, as group differences were consistently observed across *all* frequencies measured, these differences between infants’ and adults’ EEG dynamics cannot account for *specific* rate effects in the data.

### 2.9 Statistical analyses

To assess which, if any, aspects of speech-brain entrainment differed significantly between infants and mothers, the PLV indices were entered into a Repeated Measures ANOVA with Group (mother or infant) as the between-subjects factor. Hemisphere (left or right) and Frequency (7 temporal rates) were within-subjects factors. If a main effect of Group was observed, such an effect would indicate that speech-brain entrainment was, on average across all frequencies, stronger in the Infant or the Adult group. If such an effect was observed we then planned to assess interactions between Group and Frequency, to assess whether the *relative* Group difference in speech-brain entrainment varied as a function of Frequency (which was our main hypothesis). If such an interaction was observed, we planned to conduct *post hoc* Tukey HSD tests to assess at which particular frequencies Group differences were significant, or not significant (with particular attention to relative group differences at Theta and Delta rates). As secondary analyses we planned to examine: a) main effects of Frequency, to assess whether phase-locking varied significantly as a function of frequency; b) main effects of Hemisphere, to assess whether phase-locking was stronger overall from the left, or right, hemisphere, and c) Frequency x Hemisphere interactions.

## 3 RESULTS

Following the stringent, two-stage artifact rejection criteria used, 19 of the 29 infants participating in the study gave data of both sufficient quantity and quality for inclusion in the final analyses. Infants were excluded from the analyses if they contributed less than 8.0s of still, attentive and artifact-free data. For consistency, data from the parents of the infants who failed to provide enough usable data were also excluded. Therefore data from 38 participants (19 infants, 19 adults) was included in the final sample. The median age of these infants was 7.56 months (range = 6.30 to 14.82 months). On average, the 19 dyads that were retained contributed 49.0s (range 8.9s to 129.6s) of still and attentive data per pair, with each mother and infant within each pair contributing the same amount of data. Figure 3 shows the absolute phase-locking values (PLV) obtained for infants and their mothers.

### 3.1 Main Group effect

Overall we observed a marginally significant main effect of Group (F(1, 18)=4.36, p=.051, η^2^_p_ = 0.20), with infants showing stronger phase-locking than their mothers on average across all frequencies examined. However, as predicted, significant Group interaction effects emerged from the analysis revealing that relative to adults, infants showed enhanced processing at certain temporal rates but not at others (see Figure 4).

**Figure 4.**
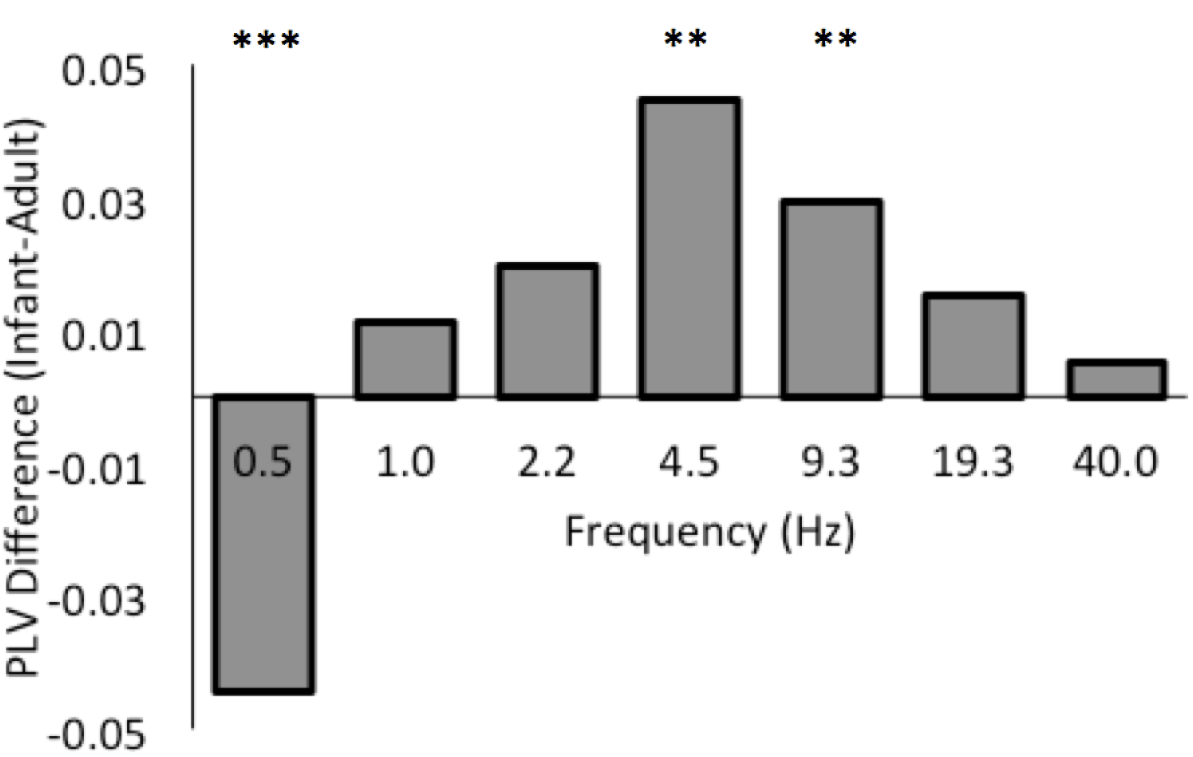
Mean group difference between infants’ and adults’ phase-locking values (infant PLV - adult PLV) averaged over left and right hemispheres, for 0.5-40 Hz. Significant group differences at 0.5 Hz, 4.5 Hz and 9.3 Hz are highlighted (i.e. these infant-adult PLV values differ significantly from 0). All differences at other rates are non-significant.

### 3.2 Group x Frequency

As illustrated in Figure 4 which plots the difference in PLV scores between infants and adults, there was a highly significant interaction between Group and Frequency (F(6,108)=18.42, p<.0001, η^2^_p_ = 0.51), suggesting that infant-adult relative entrainment differs markedly as a function of temporal rate. To further understand the pattern of these effects, a series of *post hoc* Tukey HSD tests was conducted, which indicated that infants showed significantly *stronger* phase-locking to the speech signal than their mothers at Theta (4.5Hz, p<.01) and Alpha (9.3 Hz, p<.01) rates. However, infants also showed significantly *weaker* phase-locking at the very slowest frequency of 0.5 Hz (p<.001). No significant group differences were observed at the Delta rate (p>.05). Thus, relative to adults, infants showed enhanced neural tracking of the speech signal at Theta and Alpha rates, but not at the Delta rate. Conversely, infants showed significantly less neural tracking than adults at very slow sub-Delta (~0.5 Hz) temporal rates.

### 3.3 Main Frequency Effects

We observed a strong significant main effect of Frequency (F(6, 108)=323.64, p<.0001, η^2^_p_ = 0.95). Tukey HSD post hoc analyses revealed that lower temporal rates elicited higher phase-locking than higher temporal rates and all pairwise comparisons were statistically different from each other (p<.05 for all frequency comparisons). This pattern of frequency effects indicates that speech-brain phase-locking is more robust for slower temporal rates than for faster temporal rates.

### 3.4 Main Hemisphere Effects

No significant main effect of Hemisphere was observed (F(1, 18)=1.61, p=.22, η^2^p = 0.08).

### 3.5 Frequency x Hemisphere

No significant interaction between Frequency and Hemisphere was observed (F(6, 108)=.77, p=.60, η^2^p = 0.04).

### 3.6 Supplementary Coherence Analysis

As detailed in the Supplementary materials, we performed an identical analysis of the data using an alternative measure (Wavelet Coherence) in order to assess the methodological replicability of our findings. The Coherence results agreed with those obtained using the PLV measure. Compared to the PLV, there was an even stronger main effect of Group for Coherence scores (F(1, 18)=21.49, p=<.001, η^2^p = 0.54), confirming our finding that infants showed stronger neural tracking of the speech signal than their mothers overall. Also consistent with our PLV findings, the Coherence results yielded a highly significant interaction between Group and Frequency (F(6,108)=24.63, p<.0001, η^2^p = 0.58). Tukey HSD post hoc analysis of this interaction revealed that infants showed significantly stronger coherence with the speech signal at Theta and Alpha rates (as for the PLV), and also at the Delta rate. Similar to the PLV measure, at 0.5 Hz, a trend toward lower coherence in infants relative to adults was observed but this difference did not reach significance. Thus, the Coherence index yielded the same or an even stronger pattern of results than the PLV measure. The discrepancy observed at the Delta rate is likely to reflect the fact that the Coherence measure is also influenced by the EEG power spectra (whereas the PLV measure is not), and infants’ EEG spectrum contains more power than adults’ at the Delta rate.

## 4 DISCUSSION & CONCLUSION

Multi-timescale neural oscillatory entrainment to speech temporal patterns has been proposed to be a mechanism for speech parsing in adults (Giraud & Poeppel, 2012; Ghitza, 2011; Gross et al, 2013). Here, we wished to examine *infants’* neural processing of the temporal structure of sung nursery rhymes, and to compare their profile of neural entrainment across multiple timescales with that of well-matched adults (their mothers). In particular, we were interested in whether infants’ neural response to IDS would be enhanced relative to adults’ at the Delta rate (reflecting the stronger Delta rhythms in IDS) or at the Theta rate (reflecting a tendency for Theta-oscillator parsing of speech). *A priori*, we predicted that infants’ temporal processing of IDS would be enhanced relative to adults’ at the Theta rate. To assess this, we concurrently recorded the EEG of infants and their mothers while they watched videos of recorded sung nursery rhymes. To our knowledge, this is the first study to examine and contrast infants’ and adults’ neural entrainment to natural sung infant-directed speech, using a theoretical framework that has already been well-developed for understanding adult neural processing of speech (Giraud & Poeppel, 2012; Ghitza, 2011; Gross et al, 2013).

We found that on average over all timescales measured, infants showed stronger neural entrainment than their mothers to the temporal patterns in sung IDS. Detailed analysis by timescale indicated that this group difference was driven primarily by infants’ enhanced performance at Theta and Alpha rates, which significantly exceeded adult levels of accuracy. By contrast, neural entrainment at the Delta rate was evenly-matched between adults and infants, suggesting that acuity of temporal processing at the Delta prosodic rate is similar in adults and infants (and would therefore be sufficient to support early prosodic processing). Finally, we found that infants’ entrainment accuracy fell below adult levels for the very slowest rate of 0.5 Hz. A separate analysis, reported in the Supplementary Materials, used Wavelet Coherence as an alternative method for quantifying speech-brain entrainment, and a similar pattern of results was observed for this Coherence metric.

Interestingly, we observed no significant hemisphere effects in our data. There was no overall difference in the strength of entrainment between left and right hemispheres, and also no difference between hemispheres in their relative entrainment to different temporal rates in speech. This finding was surprising because in adults, phase-locking to higher (e.g. Gamma) frequencies is generally stronger over the left hemisphere whereas slow Delta and Theta rates are typically tracked more strongly over the right hemisphere (Giraud et al, 2007). This pattern of lateralisation emerges gradually over development. For example, newborn auditory responses to phoneme (Gamma)-rate modulations are bilateral rather than left-lateralised (Telkemeyer et al, 2009), but their responses to slow modulations are right-lateralised, and this pattern is maintained until 6 months of age (Telkemeyer et al, 2011). However, hemispheric lateralisation patterns are less clear when infants are exposed to natural speech, which contains temporal patterns at multiple rates, and results may depend crucially on the specific stimuli and comparisons used. For example, using optical tomography, Pena et al (2003) reported that neonates showed enhanced left-hemisphere processing for natural IDS over backward speech or silence. However, using the same neuroimaging method, Homae et al (2006) found that 3-month-old infants showed stronger right hemisphere activation for normal as compared to pitch-flattened speech. As our study only utilised real speech stimuli rather than manipulated speech, and infants’ patterns of hemispheric lateralisation were still undergoing development, these factors could explain why we did not observe strong lateralisation effects here.

### 4.1 Theta-rate temporal processing is enhanced in infants relative to adults

Infants’ entrainment accuracy exceeded adults’ specifically at Theta and Alpha rates (~4.5 - 9 Hz). For adult-directed speech, this rate range would correspond primarily to the syllable rate of utterance (~5 Hz, Greenberg et al, 2003). However, IDS is typically produced at a slower rate than ADS (Fernald & Simon, 1984), and for our sung IDS stimuli, this rate range corresponded primarily to patterns of onset-rime units (mean rate of 4.99 Hz) and phonemes (mean rate of 7.05 Hz), as detailed earlier in Table 1. Thus, the infants in our study showed enhanced processing of sub-syllabic Theta-rate temporal patterns in speech as compared to their adult mothers, even though Theta was not the dominant rate of temporal patterning in the IDS stimuli. This finding supports the view that neural processing of speech may be best performed by Theta-rate oscillators in the cortex, whose strategic hierarchical phase-amplitude relationships with other-rate oscillators may be functionally-expedient for speech processing (Giraud & Poeppel, 2012; Ghitza, 2011).

One other learning benefit that may arise from infants’ Theta-Alpha entrainment enhancement is that these rates capture the enhanced formant and vowel structure of (slower) IDS. Infants prefer to listen to IDS over adult-directed speech (Fernald, 1985) and Zhang and colleagues found that the infant brain shows enhanced processing of IDS-like formant-exaggerated speech sounds (Zhang et al, 2011). IDS is characterised by vowel hyper-articulation (Kuhl et al, 1997), and this vowel expansion should primarily affect speech modulation patterns at sub-syllabic (here, Theta-Alpha) rates. This factor could contribute to why we observed enhanced performance for infants at sub-syllabic rates, since this exaggerated Theta-rate structure would be useful for supporting language learning in infants, but would be less useful for adults who are already language experts.

More broadly, the stronger Theta entrainment observed in infants relative to adults may shed light on the wider role that neuronal oscillations play in language acquisition. As noted in the introduction, ‘perceptual tuning’ to native phonetic categories occurs by 10-12 months of age (Werker & Tees, 1984). By 9 months, infants can identify words that either follow or violate the phonetic and phonotactic patterns of their native language (Jusczyk et al, 1993). Previous studies have already implicated oscillatory mechanisms in this process of neural commitment to native phoneme sounds (Ortiz-Mantilla et al, 2013; Bosseler et al, 2013) and also to native prosodic rhythm patterns (Pena et al, 2010). Here, we extend the previous literature by showing that infants’ neural oscillatory temporal processing of Theta-rate sub-syllabic (rime and phoneme) speech sounds is also elevated (relative to adults’) during this crucial stage of early language learning. Therefore, neural entrainment mechanisms are well-placed to support infants’ rapid acquisition of native phonetic and phonotactic statistics during their sensitive period for language learning.

### 4.2 Slow phrasal temporal processing is poorer in infants as compared to adults

In contrast with the enhanced entrainment observed in infants at Theta-Alpha rates, phase-locking to very slow sub-Delta (~0.5 Hz) temporal patterns in sung nursery rhymes was poorer in infants than adults. For our stimuli, this rate corresponded to temporal patterns spanning whole phrases such as *“The clock struck one”* (mean rate of 0.64 Hz). This result is partly surprising because infants are known to be able to process and learn about suprasegmental features of speech (such as prosodic patterning) even in utero (DeCasper & Spence, 1986; Mehler et al. 1978). Further, infants can utilise speech rhythm patterns to distinguish between languages of different rhythm typologies (Mehler et al., 1988, Nazzi et al., 2000, Ramus et al., 2000). Consistent with these behavioural studies, infants’ entrainment accuracy at the slow Delta rate (1.04 Hz), which reflected prosodic stress motifs in our stimuli (e.g. trochees and iambs), was equivalent to that of adults’.

Infants’ poorer neural tracking of larger phrasal and sentential structures cannot be explained by a lack of modulation power in the nursery rhyme stimuli at slow rates, as the modulation spectrum of the stimuli indicated that there was strong power at rates under 1 Hz. Rather, the difference between infant and adult performance could have occurred because the infants in this study are at a very early stage of syntactic development where they do not yet combine single words into phrases (Gleason, 1985), and therefore they may not prioritise tracking very slow temporal patterns in the speech signal that correspond to these larger phrasal units. Consistent with the evidence that *phonetic* learning is enhanced at this early developmental stage, one explanation could be that the strength of speech-brain entrainment reflects infants’ current stage in language development (i.e. the level of linguistic analysis that infants are able to perform). Alternatively, it is also possible that very slow oscillatory tracking mechanisms are not yet mature at this stage in infancy. As no other prior studies with infants (to our knowledge) have examined early neural tracking of acoustic patterns at such slow rates, it is not possible at this point to distinguish between these two explanations. However, a recent *adult* MEG study by Ding et al (2016) suggests that adult cortical activity does indeed track much slower and larger linguistic structures such as phrases and sentences, and that the neural tracking at these higher levels is less acoustically-driven and more dependent on grammar-based (knowledge) construction. This predicts that adults, who have greater knowledge about sentence structure, should show indeed stronger neural tracking of these very slow structures than infants who have less sentential knowledge, which is consistent with our current result.

### 4.3 Limitations

One limitation to the current study is that brain activity was measured only at two specific scalp locations (C3 and C4), in accordance with the expected fronto-central topography of ASSR signals (e.g. Saupe et al, 2009; Shuai & Elhilali, 2014). However, our pilot results indicated similar findings in infants when electrodes were placed over frontal F3 and F4 sites instead. This observation is consistent with a previous study by Wunderlich et al (2006) on the development of cortical auditory evoked potentials across newborns, toddlers, children and adults. Wunderlich et al reported that for newborns and toddlers (13-41 months), the N1, N2 and P2 components were uniformly distributed across the scalp over fronto-central and temporal regions, whereas adults showed a more focal distribution pattern with a peak around central electrodes. By contrast, there was no change in the P1 scalp distribution pattern with age, which remained uniformly distributed across the scalp in all age groups. Accordingly, by recording only from central sites in this experiment, we expect to have captured the peak adult auditory response, as well as a representative picture of a spatially evenly-distributed infant response. In future it will be useful to assess the replicability of these findings over other neural recording sites, using high-density arrays.

Although the PLV index is insensitive to the amplitude of the neural response, it may be affected by the timing (or latency) of the neural response. Adults’ neural response latencies are typically shorter than infants’ (Wunderlich et al, 2006; Oades et al, 1997; Ponton et al, 2000; Shucard et al, 1987). For example, the P1 latency to high-pitched tone stimuli decreases from a mean value of 85.7 ms in newborns and 79.1 ms in toddlers to just 56.0 ms in adults (Wunderlich et al, 2006). Thus, if the time window of analysis is not sufficiently large (e.g. 60 ms), infants’ neural response to a given section of the stimulus may be missed (because it occurs after the end of the time window of analysis) whilst the adults’ neural response is retained within the time window of analysis, producing artifactually low entrainment values for infants. Here, the time window of analysis used was 2.0s (2000 ms) which avoided this error.

Our analyses indicated that there was no significant association between infant age and the degree of speech-brain synchronisation observed in our cohort (infant mean age of 9.1 months). In future studies, it would be interesting to assess whether even younger infants show the same boost in sub-syllable rate phase-locking to speech, and at what developmental timepoint this enhanced processing emerges.

A further limitation is that audio-visual stimuli were presented, meaning that we cannot be sure whether the entrainment observed was due to the auditory or the visual aspects of the stimulus. Of note, attention to the mouth changes with age (Lewkowicz & Hansen-Tift, 2012), and infants are more sensitive to the temporal correlation between articulatory movements and speech (Baart, Vroomen, Shaw, & Bortfeld, 2014). Future work should investigate this question using audio and visual stimuli separately.

A further limitation to the study is that the IDS stimuli used, whilst high in ecological validity for the infant, are complex stimuli that may potentially differ for the adult and infant in terms of familiarity and attentional capture. Accordingly, the adult-infant differences in phase-locking observed may also reflect differences in higher-order semantic and attentional processing that moderate basic perceptual tracking of the speech envelope. To ameliorate these potential confounds, we ensured that the nursery rhymes we used were familiar for both mothers and infants, and we took care to only analyse matching attentive periods (identified through video coding) from infants and their mothers. Nonetheless, in future studies, it would be of theoretical importance to assess whether infants also show enhanced neural entrainment for non-speech stimuli that are equally unfamiliar and neutral for both infants and adults. A further constraint to the generalisability of our findings is that sung speech was used as the stimulus, rather than spoken IDS. As noted in the Methods section, the use of sung IDS meant that key phonological units like syllables and phonemes were produced on a slower timescale than for spoken IDS. Thus, to assess the generalisability of these findings, it will be necessary to conduct further analyses using adult-directed as well as non-sung and spontaneous infant-directed language, to assess the degree to which findings are specific to sung infant-directed language.

### 4.4 Conclusion

Overall our results suggest that by the end of the first year of life, infants show a high fidelity of neural entrainment to sung nursery rhymes, and their performance is enhanced relative to adults’ at Theta and Alpha rates (corresponding to sub-syllabic units in our stimuli), and but poor than adults’ at very slow (0.5 Hz) rates (corresponding to phrasal units in our stimuli). The enhancement in infants’ neural tracking of Theta-rate temporal structure may suggest that Theta-rate oscillators are best equipped to provide the high temporal acuity required by language-learning infants to accurately parse the fine-grained spectro-temporal structure of the speech signal for its phonemic and phonotatic structure during a sensitive period for phonetic learning. Future work should investigate the degree to which these differences are specific to the sung, infant-directed nursery rhymes that we used. If it can be shown that they are also present for other, adult speech sounds, these findings may cast important new light on the role that neuronal oscillations play in early language acquisition, and shed light on the developmental relationship between neural entrainment and language processing abilities.

## Footnotes

1 https://www.neurobs.com/menu_support/menu_forums/view_thread?id=9605

## ACKNOWLEDGEMENTS

This research was funded by an ESRC Transforming Social Sciences grant ES/N006461/1 to VL and SW, a Lucy Cavendish College Junior Research Fellowship to VL, and a British Academy Post-Doctoral Fellowship to SW.

## Supplementary Methods

The analyses presented in the main text use phase locking value to index speech-brain entrainment. Here we present additional analyses that use an alternative mathematical method: Wavelet coherence.

The main difference between WCOH and PLV is that WCOH reflects both the *power* and *phase* coupling between two time series, whereas the PLV measure just measures phase synchronisation. Infant EEG responses are typically larger in amplitude than adult responses (Kurtzberg et al, 1984), but this larger amplitude is interpreted as reflecting developmental immaturity rather than enhanced processing ability. Accordingly, neural entrainment indices (such as coherence) that are sensitive to both phase and amplitude provide a less ideal basis for comparing between infant and adult responses (i.e. stronger coherence may be due to stronger phase-locking and/or higher amplitude). By contrast, a pure phase-locking measure such as the PLV that is in sensitive to amplitude differences allows for a more straightforward interpretation of any observed differences between adults and infants.

### Wavelet coherence (WCOH)

quantifies the coherence between two time series as a function of both time and frequency (Torrence & Compo, 1998; Grinsted et al, 2004). This metric is well suited to investigating changes in coupling between nonstationary time series, and thus is particularly appropriate for use with neural data (Chang & Glover, 2010). WCOH utilises the continuous wavelet transform, which performs a time-frequency decomposition by convolving the time series with scaled and translated versions of a wavelet function (Mallat, 1999). Here, the wavelet function chosen was the complex Morlet wavelet (bandwidth of mother wavelet = 1 Hz, time resolution = 0.1 Hz) and the wavelet transform was computed at 7 frequencies, log-spaced between 0.5 Hz to 40 Hz. The Matlab function ‘wcoher’ (Mathworks, Inc) was used to estimate the wavelet coherence between the EEG signal and the speech amplitude envelope. WCOH values range between [0,1], and can be conceptualized as the localised correlation coefficient in time and frequency space (Grinsted et al., 2004).

### Methodological control analysis with white noise surrogates and shuffled data

To assess whether the data yielded levels of coherence that were higher than that which would be expected to occur by chance, a methodological control analysis was conducted with white noise surrogates, identical to the analysis described in section 2.7 of the main text. As with the phase-locking values, all coherence experimental data was significantly different from the white noise threshold values at all frequencies (Benjamini-Hochberg FDR-corrected p<.05). Similarly, threshold values were computed by shuffling the data in 20ms-long segments to assess the effect of destroying the temporal correspondence between the EEG and stimulus patterns. All infant data values were significantly higher than their corresponding shuffled data values (Benjamini-Hochberg FDR-corrected p<.05), but for the adult data, Beta (19 Hz) and Gamma (40 Hz) WCOH values were not significantly above shuffled data thresholds for both the left and right hemispheres. Therefore, the adult WCOH data at these higher frequencies do not reliably reflect significant coherence with the speech signal. For reference, the white noise and shuffled data thresholds are annotated in Figure S1 of the Supplementary Results.

## Supplementary Results

To assess which, if any, aspects of speech-brain entrainment differed significantly between infants and mothers, the WCOH indices were entered into a repeated measures ANOVA with Group (mother or infant) as the between-subjects factor. Hemisphere (left or right) and Frequency were within-subjects factors. In the following sections, the main Group effect and interactions between Group x Frequency and Group x Hemisphere are presented first, followed by the other Frequency and Hemisphere effects. This analysis is identical to the analyses presented for PLV in the main text.

**Figure S1.**
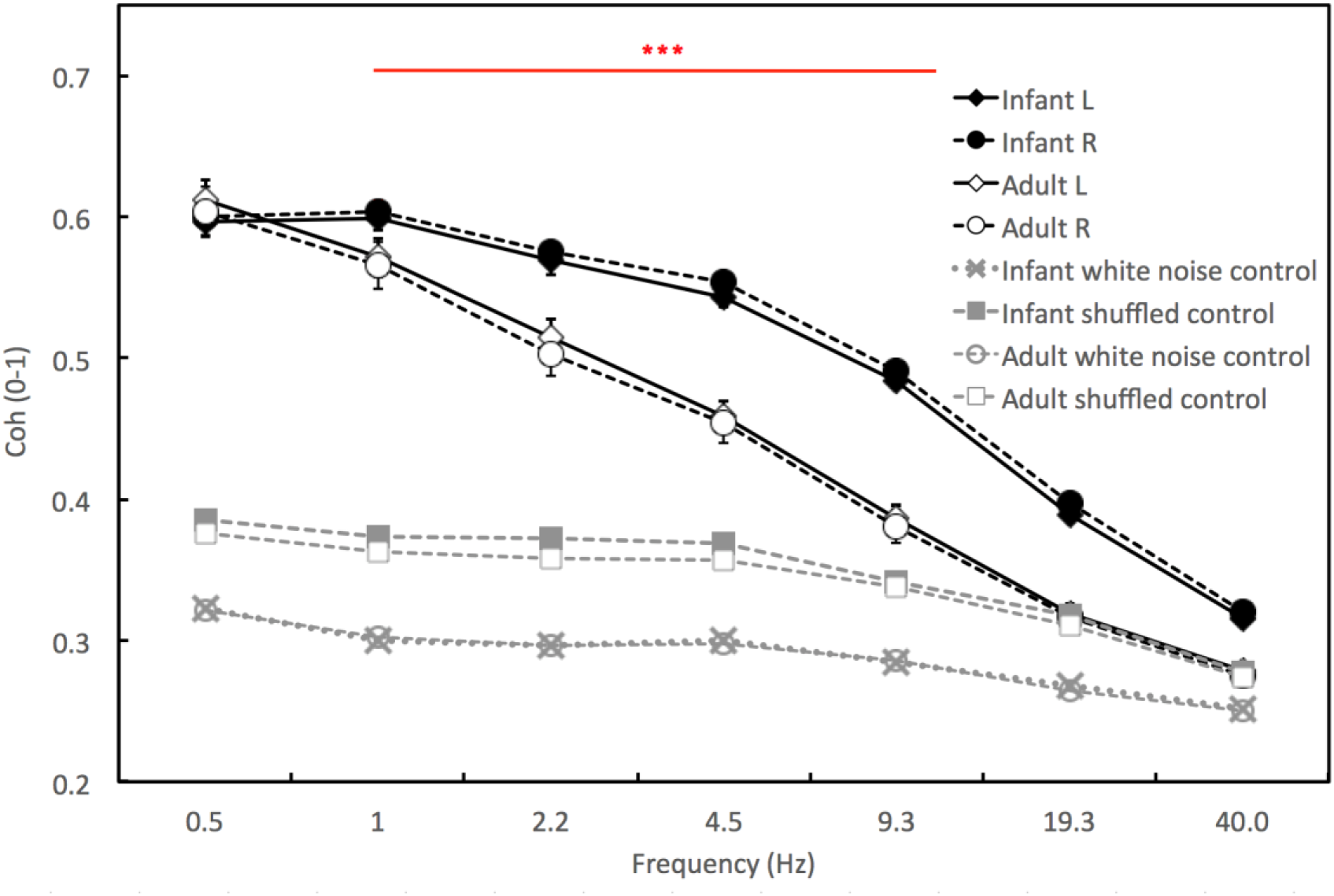
Wavelet spectral coherence (WCOH) scores for infants and adults over left and right hemispheres, for 0.5-40 Hz. ‘Infant L’ and ‘Infant R’ show the results obtained over the left and right hemisphere electrodes, accordingly. ‘White noise threshold’ show the results of the control analysis conducted with white noise surrogates, as described in the Methods section. ‘Shuffled control’ shows the results of the second control analysis conducted with shuffled data, as described in the Methods section. Error bars show Standard Error of the Means. Stars indicate the significance of the adult vs infant Tukey post hoc HSD tests, conducted as described in the main text. ** p<.01, *** p<.001. Comparisons with the control data were significant for all comparisons with the exception of the Adult L/R vs Adult shuffled control comparison at Beta (19Hz) and Gamma (40Hz).

### 1.1 Main Group effects

A highly significant effect of Group for spectral coherence scores was observed (F(1, 18)=21.49, p=<.001, η^2^_p_ = 0.54) indicating that, overall, infants showed stronger neural coherence with the speech signal as compared to their mothers. This result is stronger than the marginally non-significant (p=.051) effect reported for PLV in the main text. However, as with PLV (Figure 4) the Group interaction effects that emerged from the analysis revealed that there was a more complex pattern of developmental differences between infants and adults, with infants showing enhanced processing at certain temporal rates relative to adults.

### 1.2 Group x Frequency

There was a highly significant interaction between Group and Frequency for spectral coherence scores (F(6,108)=24.63, p<.0001, η^2^_p_ = 0.58). A *post hoc* Tukey HSD test indicated that infants showed stronger neural coherence with the speech signal at all frequencies greater than 0.5 Hz (p <.001 for frequencies including delta, theta, alpha, beta and gamma frequency ranges). At 0.5 Hz a trend effect of lower coherence in infants relative to adults was observed but the difference did not reach significance.

### 1.3 Main Frequency Effects

A strong significant main effect of Frequency was observed (F(6, 108)=891.41, p<.0001, η^2^_p_ = 0.98). Tukey HSD post hoc analyses revealed that lower temporal rates elicited higher phase-locking than higher temporal rates (i.e. a systematic downward slope), and all pairwise comparisons were statistically different from each other (p<.05 for all frequency comparisons). These results are identical to those observed for PLV, as reported in the main text.

### 1.4 Main Hemisphere Effects

No significant main effect of Hemisphere (F(1, 18)=.0008, p=.98, η^2^_p_ = 0.00). These results are identical to those observed for PLV, as reported in the main text.

### 1.5 Frequency x Hemisphere

No significant interaction between Frequency and Hemisphere was observed (F(6, 108)=.45, p=.84, η^2^_p_ = 0.02). These results are identical to those observed for PLV, as reported in the main text.

## Supplementary Discussion

### Consistency between entrainment measures

Here, we analysed neural entrainment to the speech signal using two mathematically distinct measures: coherence and phase-locking value. The pattern of results was broadly similar for both metrics (see Figure 3 and Figure S1). Both metrics were consistent in indicating that infant entrainment was enhanced relative to adults at Theta-Alpha rates. Although coherence was elevated in infants relative to adults for Beta and Gamma rates additionally, adult WOCH values at these frequencies were not significantly above the threshold established by shuffling the data to destroy internal temporal correspondences. Thus, whilst infants showed reliable coherence at Beta and Gamma rates, adults did not, and therefore a group comparison is not statistically possible. At the slowest 0.5 Hz rate, although adult coherence values were not statistically greater than infants’, there was a trend in the same direction as observed for the PLV metric. Finally, there was one important discrepancy between the PLV and WCOH results. Whilst infant PLVs were matched to adults’ at the Delta rate, their WCOH Delta values *exceeded* adults’.

As the phase-locking measure uses only the phase of the signals and not their power, this discrepancy can be explained if infants possess higher EEG power than adults at the Delta rate, resulting in stronger coherence with the speech signal at the Delta rate but not stronger phase-locking. Inspection of Figure S3, which shows the actual EEG power spectra of the participants, reveals that this is indeed the case. Infants possess higher power in their EEG spectra at Delta and Theta rates, which could explain why infants WCOH values were higher than adults’, but their PLV values were not.

**Figure S3.**
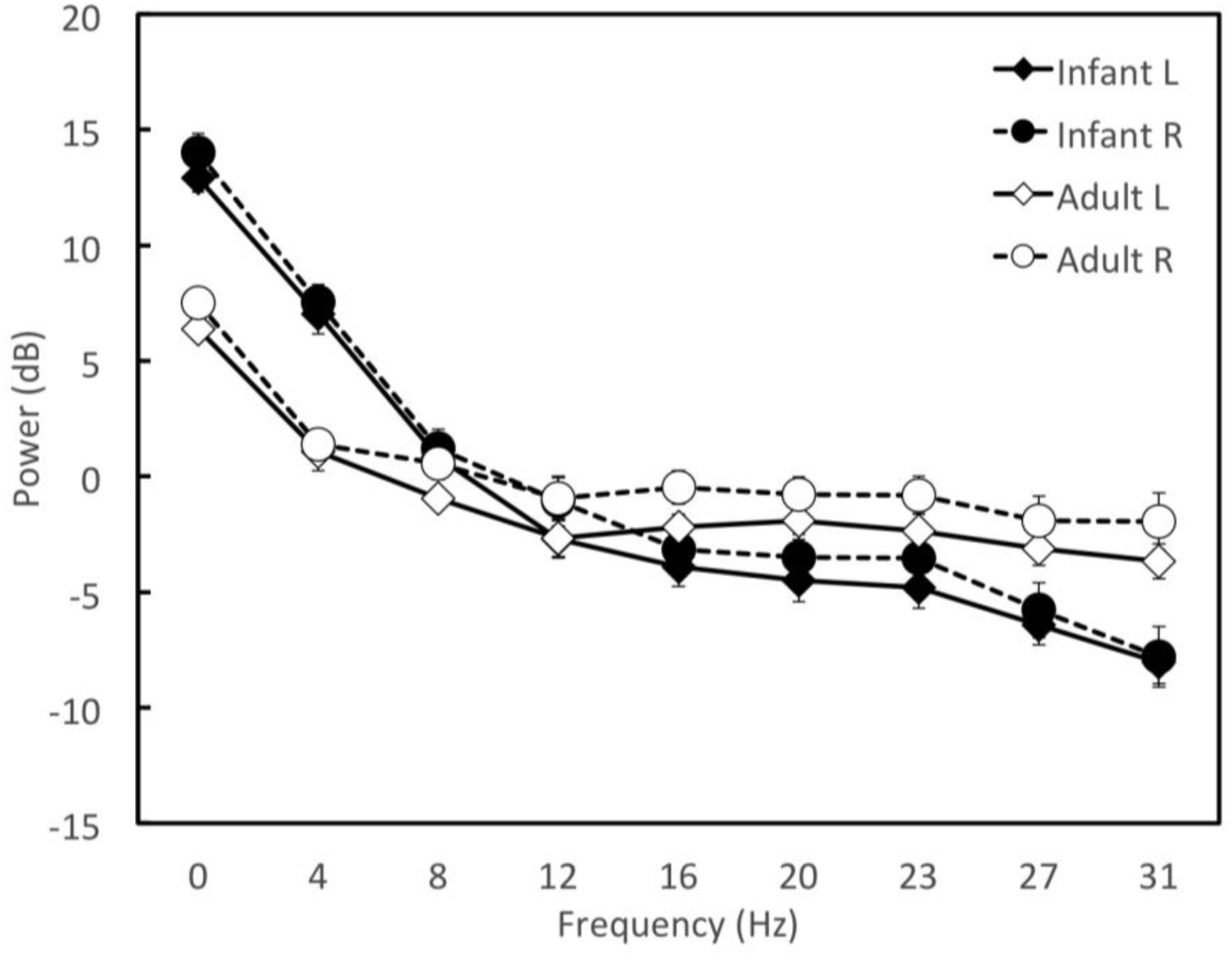
Mean EEG power for infants and adults over left (C3) and right (C4) electrodes.

